# Spatial ecology of Paraguay’s last remaining Atlantic forest jaguars (*Panthera onca*): implications for their long-term survival

**DOI:** 10.1101/426940

**Authors:** Roy T McBride, Jeffrey J Thompson

## Abstract

Using GPS telemetry we quantified space use and movements of jaguar (*Panthera onca*) in remnant populations in the Paraguayan Atlantic forest within a comparative context with populations in the Argentine and Brazilian Atlantic forest. Mean estimated home range size was 160 km^2^; estimated to be nearly equal to jaguars in the Morro do Diabo State Park in Brazil but jaguars in other populations in Argentina and Brazil had a 73% (Iguazú/Iguaçu national park complex) and 96% (Ivinhema State Park) probability of having larger home ranges. We found no relationship between home range size or movements and human population or the Human Footprint Index, while 75% of locations from all individuals were in protected areas. Our data and analysis highlight the dependence of Atlantic forest jaguars on protected areas, an avoidance of the landscape matrix and an extreme isolation of the remaining Paraguayan Atlantic forest jaguars.

## Introduction

The populations and distributions of large carnivores are in decline globally with significant implications for the altering ecosystem dynamics and function and corresponding loss of ecosystem services (Estes et al. 2011; Ripple et al. 2012). In the majority of the Neotropics, the jaguar (*Panthera onca*) is the dominant predator, however, it was been extirpated from the majority of its historic distribution due to habitat loss, prey depletion, and persecution (de la Torre et al. 2017). In South America the population reduction and range contraction of jaguar has been particularly acute in the Atlantic forest biome (De Angelo et al. 2013; Paviolo et al. 2016).

The Atlantic forest formerly occupied >1.5 million km^2^ in Argentina, Brazil and Paraguay, however, habitat conversion has greatly reduced its coverage to ~12% of its original area, >30% of which remains in fragments of <100 ha and intermediate secondary forest (Ribeiro et al. 2009). As a result of this forest loss 85% of jaguar habitat in the Atlantic forest has been lost, the remaining jaguar populations are isolated and confined to larger habitat remnants, and consequently, the long-term persistence of those populations is precarious (Paviolo et al. 2016).

In the Paraguayan Atlantic forest jaguars have been mostly extirpated, with two small populations remaining which are confined to two forest remnants in the northeast of the country; the Mbaracayu and Morombí forest reserves where <20 jaguars are present in either reserve (Paviolo et al. 2016) and these populations apparently isolated from each other and other populations in Argentina and Brazil (De Angelo et al. 2013; Thompson and Velilla 2017). Consequently, an understanding of how Paraguayan Atlantic forest jaguars utilize space, particularly the use of protected areas compared to the surrounding landscape matrix, is imperative for the conservation of those populations specifically and for Atlantic forest jaguars in general.

Although home range and movement parameters of Atlantic forest jaguars have been estimated for individuals in Argentina and Brazil (Morato et al. 2016), conspicuously no such estimates exist for jaguars in the Paraguayan Atlantic forest. Towards addressing the absence of data on jaguar spatial ecology in eastern Paraguay we estimated home range and movement parameters of Atlantic forest jaguars in the remnant populations in northeastern Paraguay using continuous time movement models and auto-correlated kernel density estimation (Fleming et al. 2014, 2015) placing those estimates in a comparative context with others from Atlantic forest jaguars.

Additionally, as jaguar home range size has been illustrated to be related to anthropogenic factors (Morato et al. 2016) we examined the relationships among anthropic and biotic factors at the landscape-scale with home range size and movement parameters of jaguars from the Atlantic forest in Argentina, Brazil and Paraguay. By placing our results within a comparative context of the conservation status of jaguars in the Atlantic forest we examine the conservation implications of the observed spatial ecology of jaguars in the Paraguayan Atlantic forest within Paraguay specifically and overall in the Atlantic forest biome.

## Material and methods

### Study area

Our research was undertaken in the Mbaracayu and Morombí Reserves in the departments of Canindeyú and Caaguazú in eastern Paraguay, two private reserves of 644 km^2^ and 270 km^2^, respectively within the Upper Paraná Atlantic forest ecoregion (Olson et al. 2001). Both reserves are forest remnants in an agricultural matrix dominated by soybean farming and small-scale agriculture (Fig. 1) and support the last reproducing populations of jaguar in the Paraguayan Atlantic Forest (Paviolo et al. 2016).

**Figure 1.**
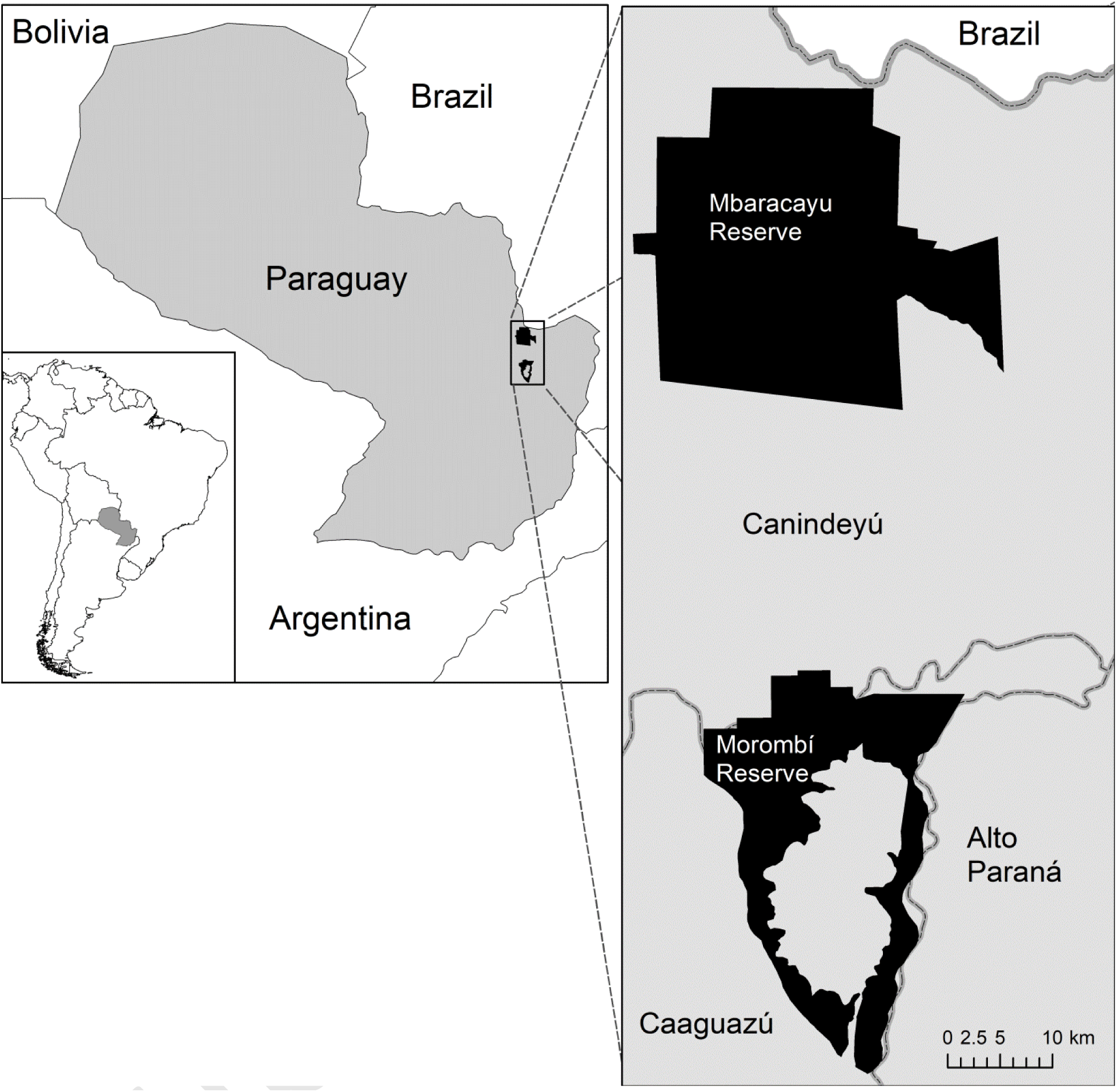
Map of location of the Mbaracayu and Morombí reserves in Paraguay.

### Data collection and analysis

Jaguars were captured in 2009 and 2010 using trained hounds to tree or bay jaguars which were then anesthetized using a weight-dependent dose of a mix of ketamine hydrochloride and xylazine hydrochloride injected by a dart shot from a tranquilizer gun (McBride and McBride 2007). Capture methods followed ASM protocols (Sikes et al. 2016). Jaguars were fitted with Northstar (D-cell, Northstar, King George, VA USA) or Teleonics (Generation 111 store-on-board, Tuscon, AZ USA) GPS collars programmed to record locations at 4 h intervals.

To estimate home ranges and movement parameters we fit continuous-time stochastic models of movement to the telemetry data, incorporating variogram analysis of semi-variance in locations in relation to time lags to inspect the autocorrelation structure in the data over time and to account for variable sampling intervals (Fleming et al. 2014). We used starting values derived from semi-variance functions for maximum likelihood model fitting, selecting best models based upon Akaike Information Criteria adjusted for small sample size (AICc) and model weights (Fleming et al. 2014, 2015; Calabrese et al. 2016).

We tested movements using a random search model (Brownian motion) with uncorrelated velocities and no limits to space use, a random search model with constrained space use (Ornstein–Uhlenbeck, OU), and Ornstein–Uhlenbeck motion with foraging (OUF) which is the OU process with correlated velocities (Fleming et al. 2014; Calabrese et al. 2016). All these models account for autocorrelation in positions, while the OUF model accounts for autocorrelation in velocities and the OU and OUF models include range residency (home range). Both the OU and OUF models produce estimates of home range size and home range crossing time, while the OUF model also estimates the velocity autocorrelation time scale (a measure of path sinuosity) and mean distance traveled per day (Fleming et al. 2014; Calabrese et al. 2016). When individuals exhibited residency in movement we estimated 95% home range areas using autocorrelated kernel density estimation (AKDE) based upon the best fitting model to account for serial autocorrelation in the data (Fleming et al. 2015). We undertook semi-variogram analysis, model selection and AKDE using the *ctmm* package (Calabrese et al. 2016) in R 3.5 (R Development Core Team 2010).

We examined the relationship of home range size and movement parameters with landscape-scale anthropogenic factors for Atlantic forest jaguars from this study and 14 Atlantic forest jaguars from Argentina and Brazil (Rio Ivinhema State Park, Mato Grosso do Sol, Brazil [3 individuals], Morro do Diabo State Park, São Paolo, Brazil [6 individuals], Iguazú National Park, Misiones, Argentina and Iguaçu National Park, Paraná, Brazil [5 individuals]; Morato et al. 2018). Using linear regression we tested the relationship between home range size and movement parameters in relation to the square root of the mean population density (*sensu* Morato et al. 2016) and the mean Human Footprint Index (HFI) value. We derived the mean human population density and HFI values from the LandScan data set (Bright et al. 2010) and the 2009 Global Terrestrial Human Footprint map (Venter et al. 2016), respectively, selecting a two decimal degree square area centered upon each home range.

We tested for differences in the estimates of jaguar home range and movement parameters from Paraguayan Atlantic forest jaguars with other jaguar populations in the Argentine and Brazilian Atlantic forest (Morato et al. 2018), accounting for sex-based differences, using a two-way analysis of variance (ANOVA) in a Bayesian framework. Home range and movement parameters were log transformed to facilitate model convergence. The analysis was undertaken using R 3.5.0 (R Development Core Team 2010) using WinBUGS (Lunn et al. 2000) and the R2WinBUGS package (Sturtz et al. 2005), running WinBugs with 3 chains of 50,000 iterations, discarding the first 5,000 iterations as a burn-in period. Convergence was confirmed by assuring that the scale reduction factor was <1.1 and through visual inspection of trace plots for lack of autocorrelation (Gelman et al. 2004).

We derived the probability of difference among estimated home range and movement parameters by sex and site by taking 100,000 random samples from the posterior distributions from the ANOVA and deriving the proportional frequency (probability, *pr*) that a value from a distribution was greater or less than the value from the posterior distribution under comparison. A frequency of 0.5 indicated no difference, while values approaching 0 or 1 indicated high probability of difference between values.

## Results

We captured and collared 4 jaguars during 2008-2011; 1 female in the Morombí Reserve and 1 male and two females in the Mbaracuyu Reserve, estimated to be between 2 and 9 years old (Table1). Collars operated between 150 and 540 days, collecting 174 to798 locations. All locations of jaguars captured in the Mbaracuyu Reserve were confined to the reserve and 83% of locations the jaguar captured in the Morombí Reserve occurred within that reserve (total mean = 96%).

**Table 1.**
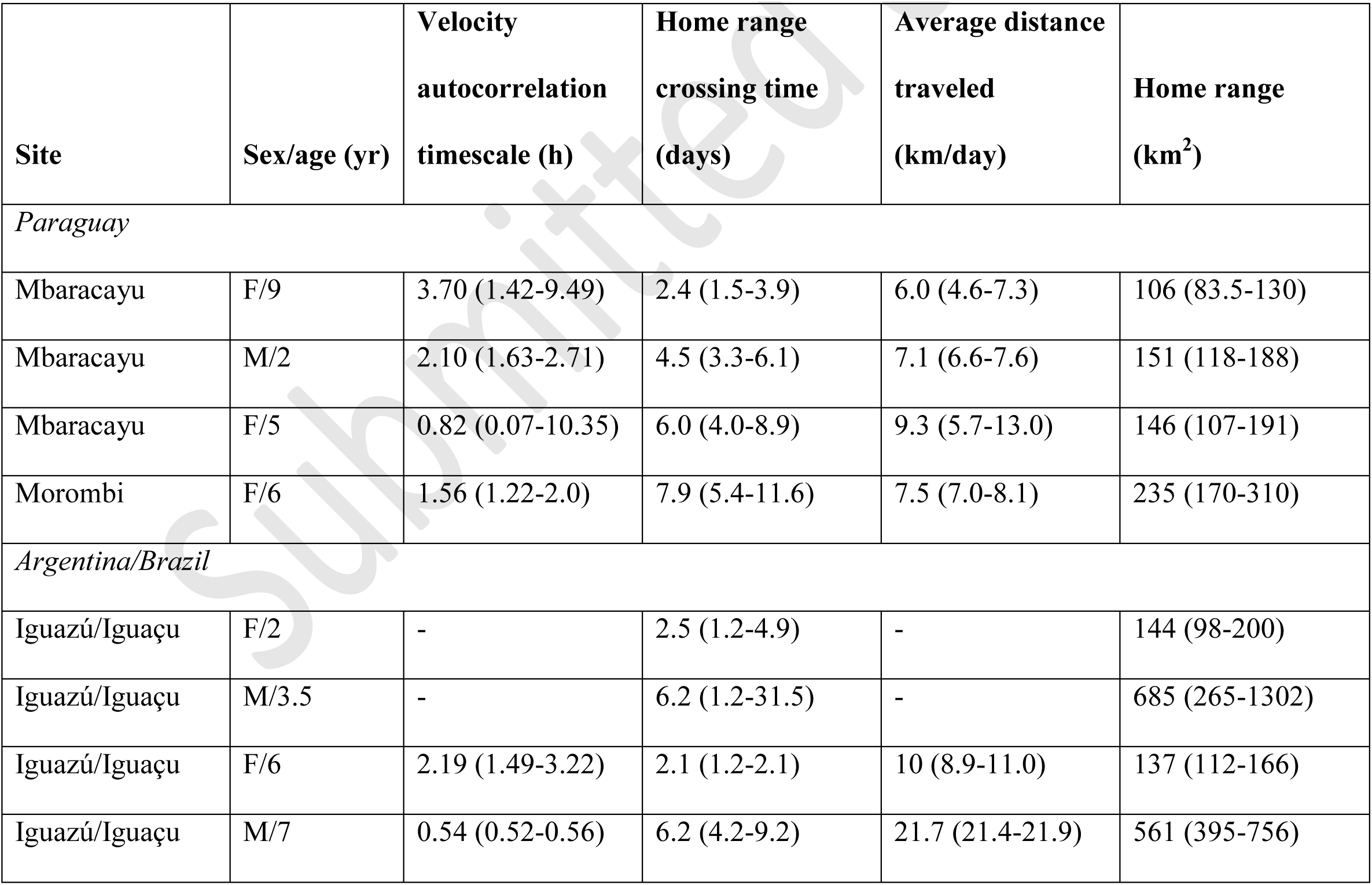
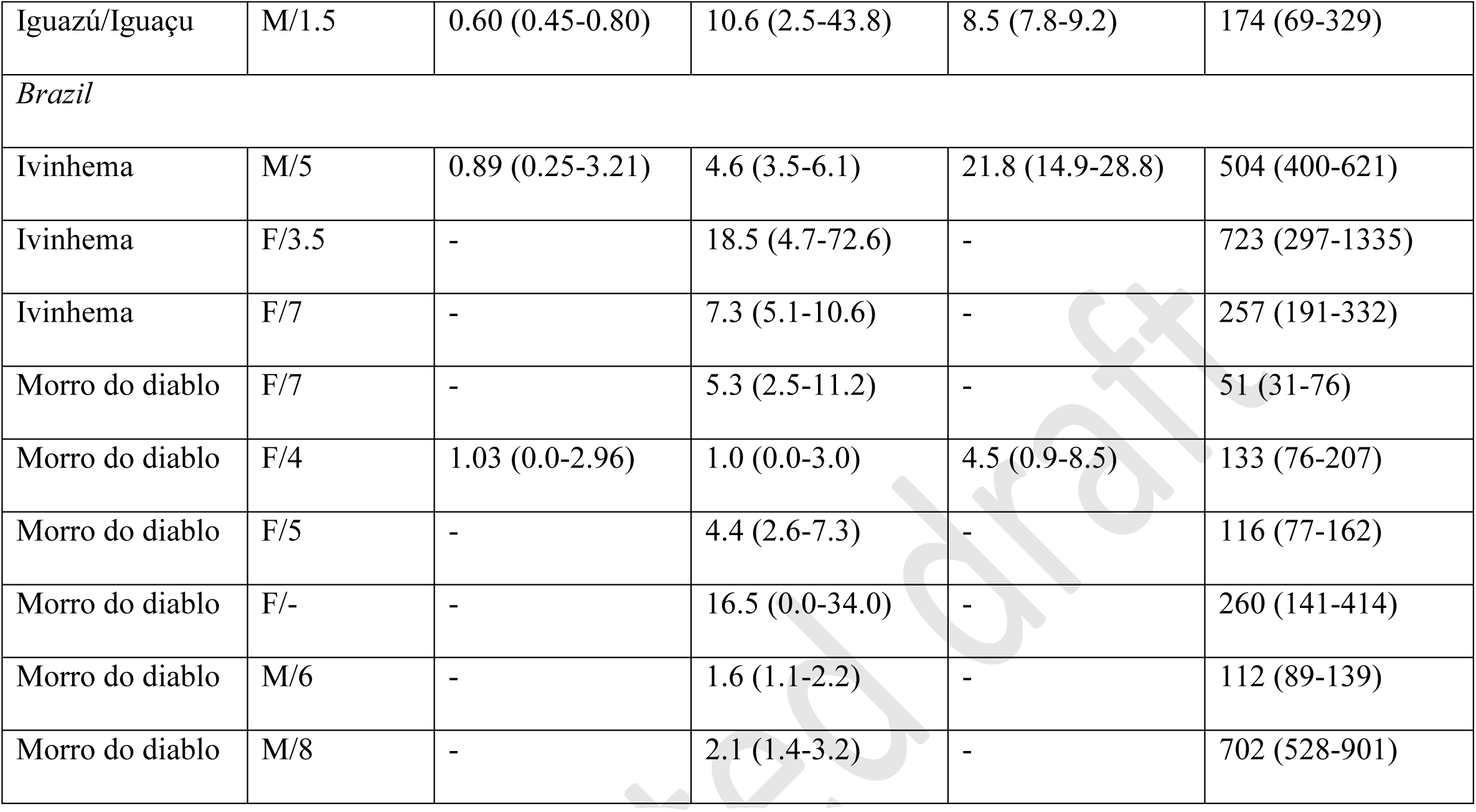
Autocorrelated kernel density home range and movement parameter estimates for Atlantic forest jaguars. In parenthesis are shown the 95% confidence intervals.

| **Site** | **Sex/age (yr)** | **Velocity**<br>**autocorrelation**<br>**timescale (h)** | **Home range**<br>**crossing time**<br>**(days)** | **Average distance**<br>**traveled**<br>**(km/day)** | **Home range**<br>**(km<sup>2</sup>)** |
| --- | --- | --- | --- | --- | --- |
| *Paraguay* |  |  |  |  |  |
| Mbaracayu | F/9 | 3.70 (1.42-9.49) | 2.4 (1.5-3.9) | 6.0 (4.6-7.3) | 106 (83.5-130) |
| Mbaracayu | M/2 | 2.10 (1.63-2.71) | 4.5 (3.3-6.1) | 7.1 (6.6-7.6) | 151 (118-188) |
| Mbaracayu | F/5 | 0.82 (0.07-10.35) | 6.0 (4.0-8.9) | 9.3 (5.7-13.0) | 146 (107-191) |
| Morombi | F/6 | 1.56 (1.22-2.0) | 7.9 (5.4-11.6) | 7.5 (7.0-8.1) | 235 (170-310) |
| *Argentina/Brazil* |  |  |  |  |  |
| Iguazú/Iguaçu | F/2 | - | 2.5 (1.2-4.9) | - | 144 (98-200) |
| Iguazú/Iguaçu | M/3.5 | - | 6.2 (1.2-31.5) | - | 685 (265-1302) |
| Iguazú/Iguaçu | F/6 | 2.19 (1.49-3.22) | 2.1 (1.2-2.1) | 10 (8.9-11.0) | 137 (112-166) |
| Iguazú/Iguaçu | M/7 | 0.54 (0.52-0.56) | 6.2 (4.2-9.2) | 21.7 (21.4-21.9) | 561 (395-756) |
| Iguazú/Iguaçu | M/1.5 | 0.60 (0.45-0.80) | 10.6 (2.5-43.8) | 8.5 (7.8-9.2) | 174 (69-329) |
| *Brazil* |  |  |  |  |  |
| Ivinhema | M/5 | 0.89 (0.25-3.21) | 4.6 (3.5-6.1) | 21.8 (14.9-28.8) | 504 (400-621) |
| Ivinhema | F/3.5 | - | 18.5 (4.7-72.6) | - | 723 (297-1335) |
| Ivinhema | F/7 | - | 7.3 (5.1-10.6) | - | 257 (191-332) |
| Morro do diablo | F/7 | - | 5.3 (2.5-11.2) | - | 51 (31-76) |
| Morro do diablo | F/4 | 1.03 (0.0-2.96) | 1.0 (0.0-3.0) | 4.5 (0.9-8.5) | 133 (76-207) |
| Morro do diablo | F/5 | - | 4.4 (2.6-7.3) | - | 116 (77-162) |
| Morro do diablo | F/- | - | 16.5 (0.0-34.0) | - | 260 (141-414) |
| Morro do diablo | M/6 | - | 1.6 (1.1-2.2) | - | 112 (89-139) |
| Morro do diablo | M/8 | - | 2.1 (1.4-3.2) | - | 702 (528-901) |

All individuals demonstrated residency with the OUF model the best fitting movement model. Mean estimated home range size was 160 km^2^ (range: 106-235 km^2^), with the mean estimated home range of females 162 km^2^ and the lone male 151 km^2^ (Table 1). Daily distance traveled was consistent among individuals (mean = 7.5 km/day; range 6.0-9.3). Home range crossing time was more variable ranging between 2.4 and 8 days, as was sinuosity (range: 0.8-3.7 hours).

Estimates of 95% home range size for jaguars from the Argentine and Brazilian Atlantic forest ranged from 51-723 km^2^ with all individuals demonstrating range residency; the majority of individuals’ (*n* = 9) movement best explained under the OU model and the others (*n* = 5) under the OUF model (Table 1). Males had a high probability of larger home ranges than females (*pr* = 0.88).

Mean estimated home range size of individuals from Paraguay demonstrated a nearly equal probability of being of similar size to those of jaguars from in and around Morro do Diabo State Park (*pr* = 0.51), with high probabilities that home range sizes were larger for jaguars associated with the Iguazú/Iguaçu National Parks (*pr* = 0.73) and Ivinhema State Park (*pr* = 0.96) compared to those from Paraguay. Home ranges from Ivinhema State Park also had a high probability of being larger than those from jaguars from Morro do Diabo (*pr* = 0.97) and Iguazú/Iguaçu (*pr* = 0.88).

Since the movement of jaguars from Argentina and Brazil mostly conformed to the OU model the autocorrelation movement factor and mean daily movement were not estimable for the majority of those individuals (Table 1) and consequently the only movement parameter that we were able to analyze comparatively by group was the home range crossing time. Home range crossing times between sexes was similar, although slightly higher for females (*pr* = 0.56). Home range crossing times for Paraguayan jaguars were greater than those in Morro do Diablo (*pr* = 0.72), equal to those from Iguazú/Iguaçu (*pr* = 0.5) and less than those from Ivinhema (*pr* = 0.8) with jaguars from Ivinhema having greater home range crossing times compared to all other sites (Table 1).

Based upon linear regression there was no relationship between estimated home range size and the square root of the mean human population density (people/km^2^; *p* = 0.83) or with the mean Human Footprint Index (*p* = 0.36). Similarly, there was no relationship between home range crossing time and the square root of the mean human population density (*p* = 0.63) or with the mean Human Footprint Index (*p* = 0.23).

## Discussion

The near extirpation of jaguar in the Paraguayan Atlantic forest has mostly confined remaining individuals to protected areas, a pattern typical of jaguar in the Atlantic forest on the whole (Paviolo et al. 2016). As was expected based upon other populations (Morato et al. 2016, McBride and Thompson 2018) male home ranges had a higher probability of being larger than those of females. Home range sizes of the Paraguayan jaguars were most similar to those associated with forest dominated protected areas in Argentina and Brazil, particularly in Morro do Diabo State Park. The larger home ranges in Ivinhema are likely associated with to a relatively small amount of forested areas and the dominant use of wetlands by jaguars at the site.

Given the degree of anthropogenic disturbance associated with the study sites in Paraguay and those in Argentina and Brazil used for comparison, as well as the documented relationship between jaguar home range size and human population, we expected to find relationships between human population and the Human Footprint Index in relation to jaguar home range size. However, the development of the Atlantic forest in the three countries share commonalties that have resulted in similar patterns of land use, infrastructure development and demographics so that there is relatively little difference among sites in regard to anthropogenic variables which may explain the lack of an observed relationship.

Moreover, all jaguar home ranges completely or partially included protected areas, with the majority of all locations within protected areas (mean = 75%), which may further explain the lack of relationship between home range size and anthropogenic factors as protected areas likely buffer negative effects of the surrounding anthropogenic matrix. Concurrently, jaguars appear to be confined to preferred habitats within or adjacent to protected areas which is apparent in the case of Atlantic forest jaguars in Paraguay where all movements of jaguars captured in the Mbaracayu reserve were confined to that reserve and 83% of locations for the jaguar from the Morombí reserve were in the reserve and remaining locations mostly located in large forest patches adjacent to the reserve.

Similarly, jaguars in the other forested protected areas in Argentine and Brazilian Atlantic forest (Iguazú/Iguaçu and Morro do Diabo) had the majority of locations occurring within protected areas (78% and 51% for Iguazú/Iguaçu and Morro do Diabo, respectively), which further supports the occurrence of a protected area boundary effect. Additional support for a boundary effect between protected areas and the surrounding matrix in affecting jaguar spatial ecology in the Atlantic forest was that the smallest home range sizes, those from Paraguay and Morro do Diabo, were associated with the smallest protected areas (Mbaracayu 664 km^2^, Morombí 270 km^2^, Morro do Diabo 339 km^2^).

The extensive loss of Atlantic forest has been the driving factor in the near extirpation of the jaguar within the biome (Paviolo et al 2016), while the remaining populations are increasingly isolated (DeAngelo et al. 2013; Paviolo et al. 2016; Thompson and Velilla 2017) and exhibit drift-induced loss of genetic diversity (Haag et al. 2010; Roques et al. 2016; Srbek-Araujo et al. 2018). The high fidelity of jaguars to protected areas and avoidance of the surrounding anthropogenic matrix suggests that exchange among the remaining populations of Atlantic forest jaguars is limited. Specifically for the Paraguayan Atlantic forest jaguars this is of concern as there is apparently little or no exchange of individuals between the Mbaracayu and Morombí reserves or with Argentine and Brazilian populations.

Apart from concerns over genetic erosion the remaining jaguars in the Paraguayan Atlantic forest are threatened by direct human impact from illegal logging, land invasion, poaching and poaching of prey species. For example, two of the four collared jaguars in this study were killed within the protected areas. Given the occurrence of poaching of jaguars and their prey within the reserves it can be inferred that the threats of low prey availability and persecution are considerably greater outside the reserves and are significant factors in limiting the successful dispersion of jaguars from the reserves.

When placed into the context of a regional landscape resistant to dispersion (DeAngelo et al. 2013; Paviolo et al. 2016; Thompson and Velilla 2017) and the observed loss of genetic diversity across the Atlantic forest, the spatial ecology of jaguars in the Paraguayan Atlantic forest indicate that the small remnant populations are isolated, dependent upon protected areas and consequently threatened by the loss of genetic diversity. Additionally, anthropogenic pressures within protected areas threaten the few remaining individuals. Consequently, there is an important need to estimate the population of jaguars and assess their genetic diversity in the Mbaracayu and Morombí reserves, reduce human impacts within the reserves and to explore mechanisms to improve connectivity between the reserves and other jaguar populations in the Atlantic forest.

## Acknowledgements

JJT received support from the Consejo Nacional de Ciencia y Tecnología - CONACYT with resources from the FEE. We thank Ronaldo Morato for comments on the manuscript

## Literature Cited

Bright, E.A., P.R. Coleman, A.N. Rose, M.L. Urban. 2010. LandScan 2009. Oakridge National Lab Oakridge, TN, USA. https://landscan.ornl.gov/

Calabrese, J.M., C.H. Fleming and E. Gurarie. 2016. “Ctmm: an R package for analyzing animal relocation data as a continuous-time stochastic process”. Methods in Ecology and Evolution. 7(9): 1124–1132.

De Angelo, C., A. Paviolo, T. Wiegand, R. Kanagaraj, and M.S. Di Bitetti. 2013. “Understanding species persistence for defining conservation actions: a management landscape for jaguars in the Atlantic Forest”. Biological conservation 159: 422–433.

Estes, J.A., J. Terborgh, J.S. Brashares, M.E. Power, J. Berger, W.J. Bond, S.R. Carpenter et al. 2011. “Trophic downgrading of planet Earth”. Science 333(6040):301–306.

Gelman, A., J.B. Carlin, H.S. Stern and D.B. Rubin, 2004. Bayesian Data Analysis. Second ed. CRC/Chapman & Hall, Boca Raton, FL.

Haag, T., A. S. Santos, D. A. Sana, R. G. Morato, L. Cullen Jr, P. G. Crawshaw Jr, C. De Angelo, M. S. Di Bitetti, F. M. Salzano, and E. Eizirik. 2010. “The effect of habitat fragmentation on the genetic structure of a top predator: loss of diversity and high differentiation among remnant populations of Atlantic Forest jaguars (Panthera onca)”. Molecular Ecology 19(22): 4906–4921.

Lunn, D.J., A. Thomas, N. Best and D. Spiegelhalter. 2000. “WinBUGS – a Bayesian modelling framework: concepts, structure, and extensibility”. Statistical Computing 10(4): 325–337.

McBride Jr., R.T. and R.T. McBride. 2007. “Safe and selective capture technique for jaguars in the Paraguayan Chaco”. The Southwestern Naturalist 52(4): 570–577.

McBride Jr., R.T. and J.J. Thompson. 2018. “Space use and movement of jaguar (Panthera onca) in western Paraguay”. Mammalia, 0(0). doi:10.1515/mammalia-2017-0040

Morato, R.G., J.J. Thompson, A. Paviolo, J.A. de La Torre, F. Lima, R.T. McBride Jr, R.C. Paula et al. 2018. “Jaguar movement database: a GPS-based movement dataset of an apex predator in the Neotropics:. Ecology 99(7): 1691.

Olson, D.M., E. Dinerstein, E.D. Wikramanayake, N.D. Burgess, G.V. Powell, E.C. Underwood, J.A. D’amico et al. 2001. “Terrestrial Ecoregions of the World: A New Map of Life on Earth A new global map of terrestrial ecoregions provides an innovative tool for conserving biodiversity”. BioScience 51(11): 933–938.

Paviolo, A., C. De Angelo, K.M. Ferraz, R.G Morato, J. Martinez Pardo, A.C. Srbek-Araujo, B. de Mello Beisiegel et al. 2016. “A biodiversity hotspot losing its top predator: The challenge of jaguar conservation in the Atlantic Forest of South America”. Scientific Reports 6: 37147

R Development Core Team, 2010. R:A language and environment for statistical computing. R Foundation for Statistical Computing, Vienna, Austria.

Ribeiro, M.C., J.P. Metzger, A.C. Martensen, F.J. Ponzoni, M.M. Hirota. 2009. “The Brazilian Atlantic Forest: how much is left, and how is the remaining forest distributed? Implications for Conservation”. Biological Conservation 142(6): 1141–1153.

Ripple, W.J., J.A. Estes, R.L. Beschta, C.C. Wilmers, E.G. Ritchie, M. Hebblewhite, J. Berger et al. 2014. “Status and ecological effects of the world’s largest carnivores”. Science. Jan 10;343(6167):1241484. doi: 10.1126/science.1241484.

Roques, S., R. Sollman, A. Jácomo, N. Tôrres, L. Silveira, C. Chávez, C. Keller et al. 2016. “Effects of habitat deterioration on the population genetics and conservation of the jaguar”. Conservation Genetics 17(1): 125–139.

Sikes, R.S. and Animal Care and Use Committee of the American Society of Mammalogists, 2016. “2016 Guidelines of the American Society of Mammalogists for the use of wild mammals in research and education”. Journal of Mammalogy 97(3): 663–688.

Srbek-Araujo, A.C., T. Haag, A.G. Chiarello, F.M. Salzano and E. Eizirik. 2018. Worrisome isolation: noninvasive genetic analyses shed light on the critical status of a remnant jaguar population. Journal of Mammalogy 99(2): 397–407.

Sturtz, S., U. Ligges and A. Gelman. 2005. “R2WinBUGS: A Package for Running WinBUGS from R”. Journal of Statistical Software 12(3): 1–16.

Venter, O., E.W. Sanderson, A. Magrach, J.R. Allan, J. Beher, K.R. Jones, H.P. Possingham et al. 2016. “Sixteen years of change in the global terrestrial human footprint and implications for biodiversity conservation”. Nature Communications 7: 12558.

